# Nanopore-based long-read transcriptome data of *Nosema ceranae*-infected and un-infected western honeybee workers’ midguts

**DOI:** 10.1101/2020.03.21.001958

**Authors:** Huazhi Chen, Xiaoxue Fan, Yu Du, Yuanchan Fan, Jie Wang, Haibin Jiang, Cuiling Xiong, Yanzhen Zheng, Dafu Chen, Rui Guo

## Abstract

*Apis mellifera ligustica* is a subspecies of western honeybee, *Apis mellifera. Nosema ceranae* is known to cause bee microspodiosis, which seriously affects bee survival and colony productivity. In this article, Nanopore long-read sequencing was used to sequence *N. ceranae*-infected and un-infected midguts of *A. m. ligustica* workers at 7 d and 10 d post inoculation (dpi). In total, 5942745, 6664923, 7100161 and 6506665 raw reads were respectively yielded from AmT1, AmT2, AmCK1 and AmCK2, with average lengths of 1148, 1196, 1178 and 1201 bp, and N50 of 1328, 1394, 1347 and 1388 bp. The length distribution of raw reads from AmT1, AmT2, AmCK1 and AmCK2 was ranged from 1 kb to more than 10 kb. Additionally, the distribution of quality score of raw reads from AmT1 and AmT2 was among Q6∼Q12, while that from AmCK1 and AmCK2 was among Q6∼Q16. Further, 5745048, 6416987, 6928170, 6353066 clean reads were respectively gained from AmT1, AmT2, AmCK1 and AmCK2, and among them 4172542, 4638289, 5068270 and 4857960 were identified as being full-length. After removing redundant reads, the length distribution of remaining full-length transcripts was among 1 kb∼8 kb, with the most abundant length of 2 kb. The long-read transcriptome data reported here contributes to a deeper understanding of the molecular regulating *N. ceranae*-response of *A. m. ligustica* and host-fungal parasite interaction during microsporidiosis.

## Value of the data

Our data presented here contributes to improving genome and transcriptome annotations of *Apis mellifera*.

This dataset can be used to identify alternative splicing and polyadenylation of mRNAs involved in response of western honeybee worker responding to *Nosema ceranae* invasion.

The accessible data facilitates better understanding the molecular mechanism underlying western honeybee response to microsporidian and host-parasite interaction during microsporidiosis.

## Data

The shared long-read transcriptome data were produced from Oxford Nanopore sequencing of *N. ceranae*-infected midguts (AmT1, AmT2) and un-infected midguts (AmCK1, AmCK2) of *A. m. ligustica* workers at 7 d and 10 d post inoculation (dpi). In total, 5942745, 6664923, 7100161 and 6506665 raw reads were yielded from AmT1, AmT2, AmCK1 and AmCK2, respectively (**Table 1**); the average lengths were 1148, 1196, 1178 and 1201 bp, and the average N50 were 1328, 1394, 1347 and 1388 bp, respectively (**Table 1**). As **Figure 1** presented, the length of raw reads from AmT1, AmT2, AmCK1 and AmCK2 was ranged from 1 kb to more than 10 kb, with the highest percentage of 1 kb. Additionally, the distribution of quality (Q) score of raw reads from AmT1 and AmT2 was among Q6∼Q12, while that from AmCK1 and AmCK2 was among Q6∼Q16 (**Figure 2**); the most abundant Q score of the above-mentioned four groups was Q9 (**Figure 2**). As **Table 2** shown, 5745048, 6416987, 6928170, 6353066 clean reads were respectively gained from AmT1, AmT2, AmCK1 and AmCK2, and among them 4172542 (72.63%), 4638289 (72.28%), 5068270 (73.15%) and 4857960 (76.47%) were identified as full-length clean reads, with the length distribution ranging from 1 kb to more than 10 kb (**Figure 3**). Moreover, after removing redundant reads, the length distribution of remaining full-length transcripts was among 1 kb∼8 kb, with the most abundant length of 2 kb (**Figure 4**).

**Table 1.** Overview of raw data derived from Nanopore sequencing

| Sample | Raw reads number | Base number | N50 (bp) | Mean length (bp) | Maximum length (bp) |
| --- | --- | --- | --- | --- | --- |
| AmT1 | 5942745 | 6822570594 | 1328 | 1148 | 14890 |
| AmT2 | 6664923 | 7976689232 | 1394 | 1196 | 16430 |
| AmCK1 | 7100161 | 8368331508 | 1347 | 1178 | 13936 |
| AmCK2 | 6506665 | 7816378025 | 1388 | 1201 | 31074 |

**Table 2.**
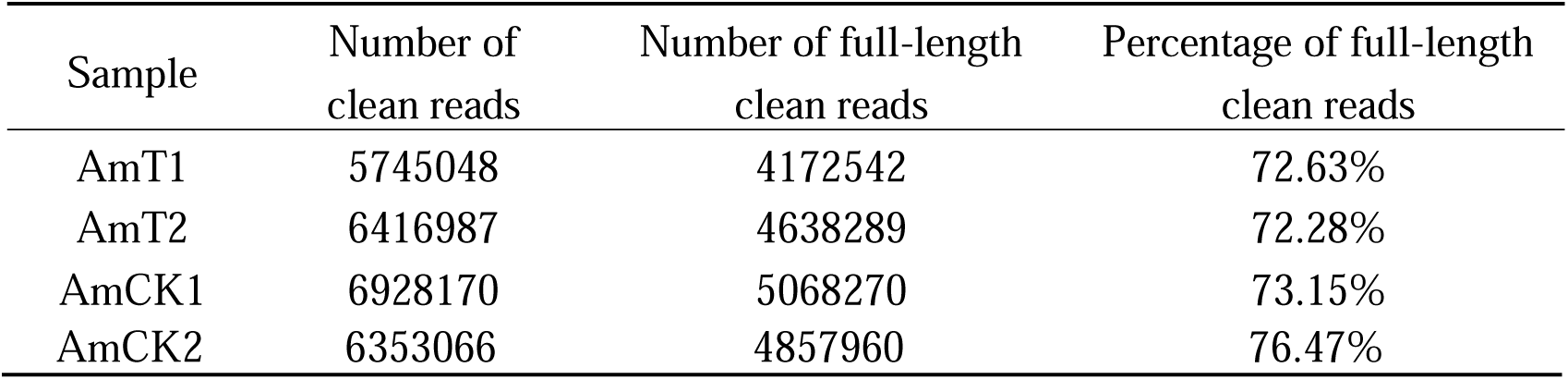
Summary of full-length clean data

**Figure 1.**
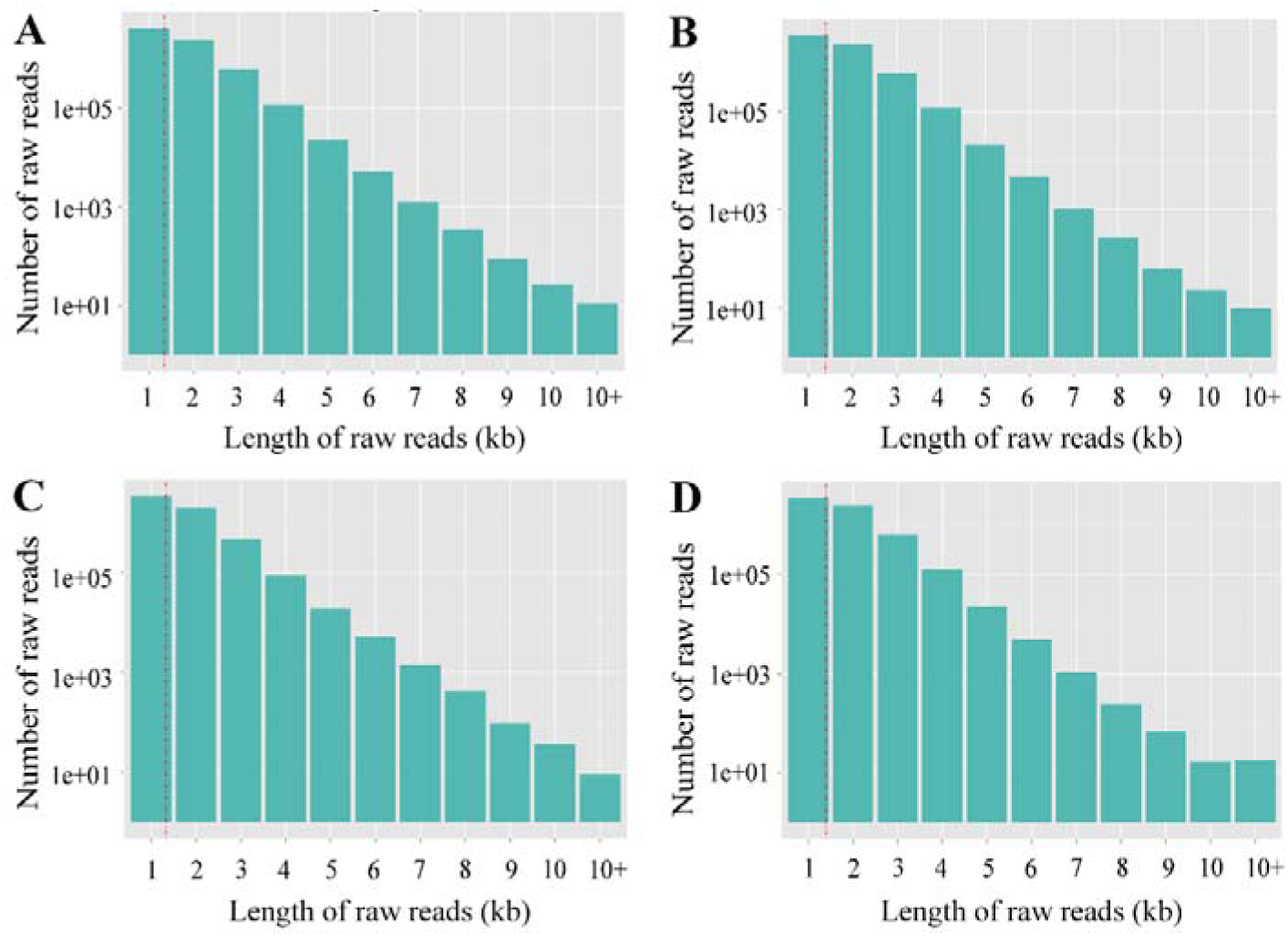
Length distribution of raw reads produced from Nanopore long-read sequencing. (A) Raw reads from AmCK1; (B) Raw reads from AmCK2; (C) Raw reads from AmT1; (D) Raw reads from AmT2.

**Figure 2.**
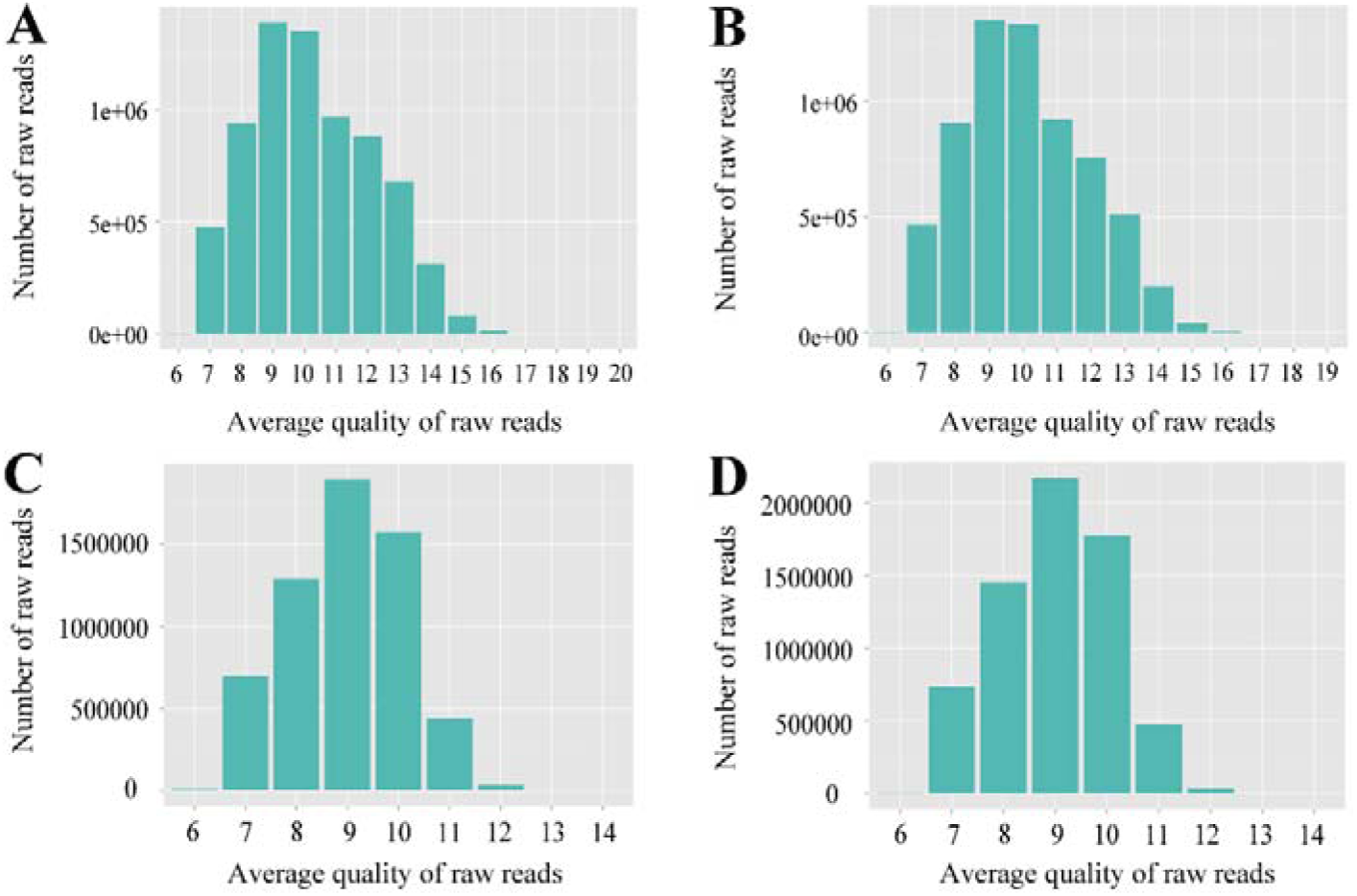
Quality distribution of raw reads produced from Nanopore long-read sequencing. (A) Raw reads from AmCK1; (B) Raw reads from AmCK2; (C) Raw reads from AmT1; (D) Raw reads from AmT2.

**Figure 3.**
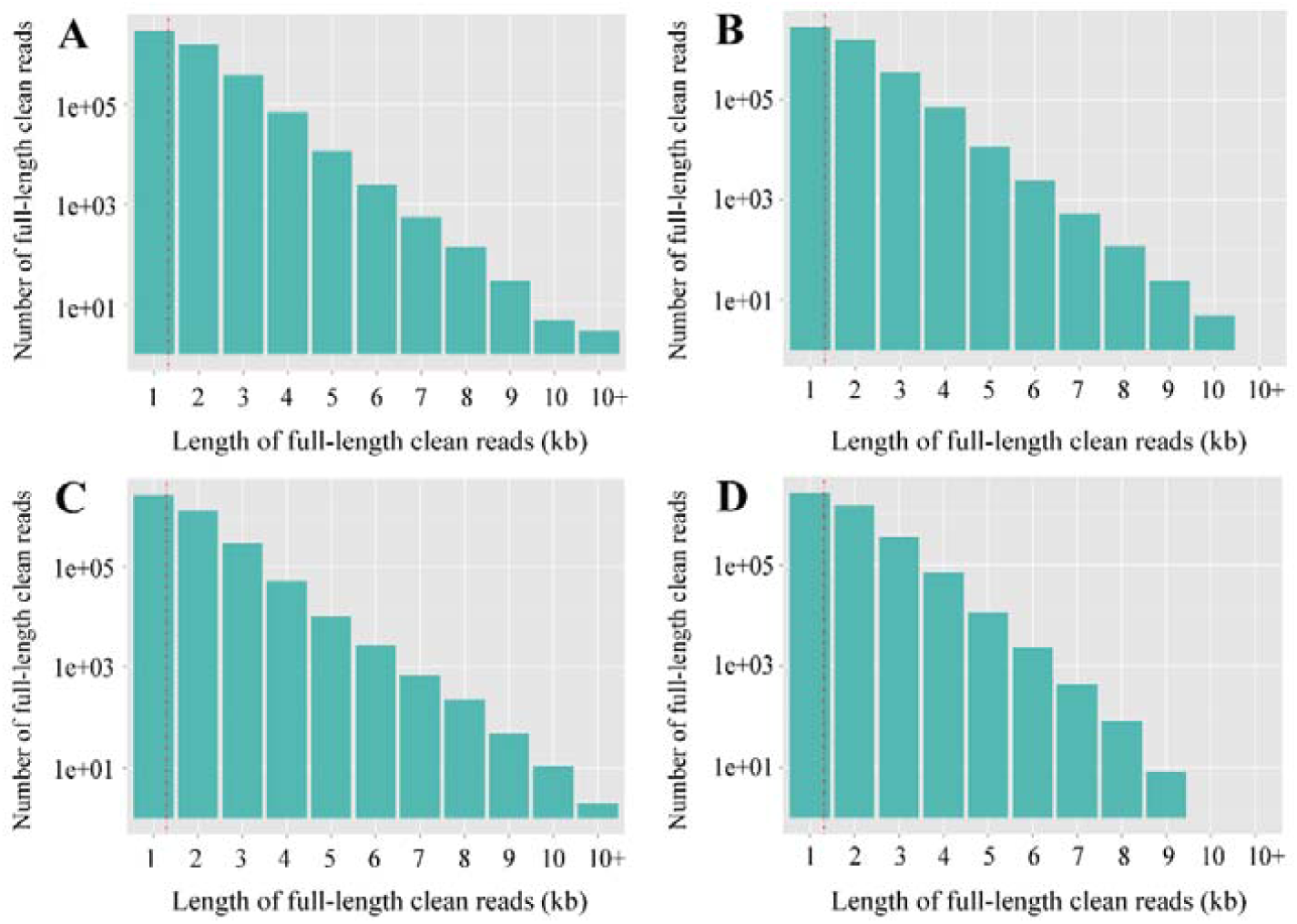
Length distribution of full-length clean reads from AmCK1 (A), AmCK2 (B), AmT1 (C) and AmT2 (D).

**Figure 4.**
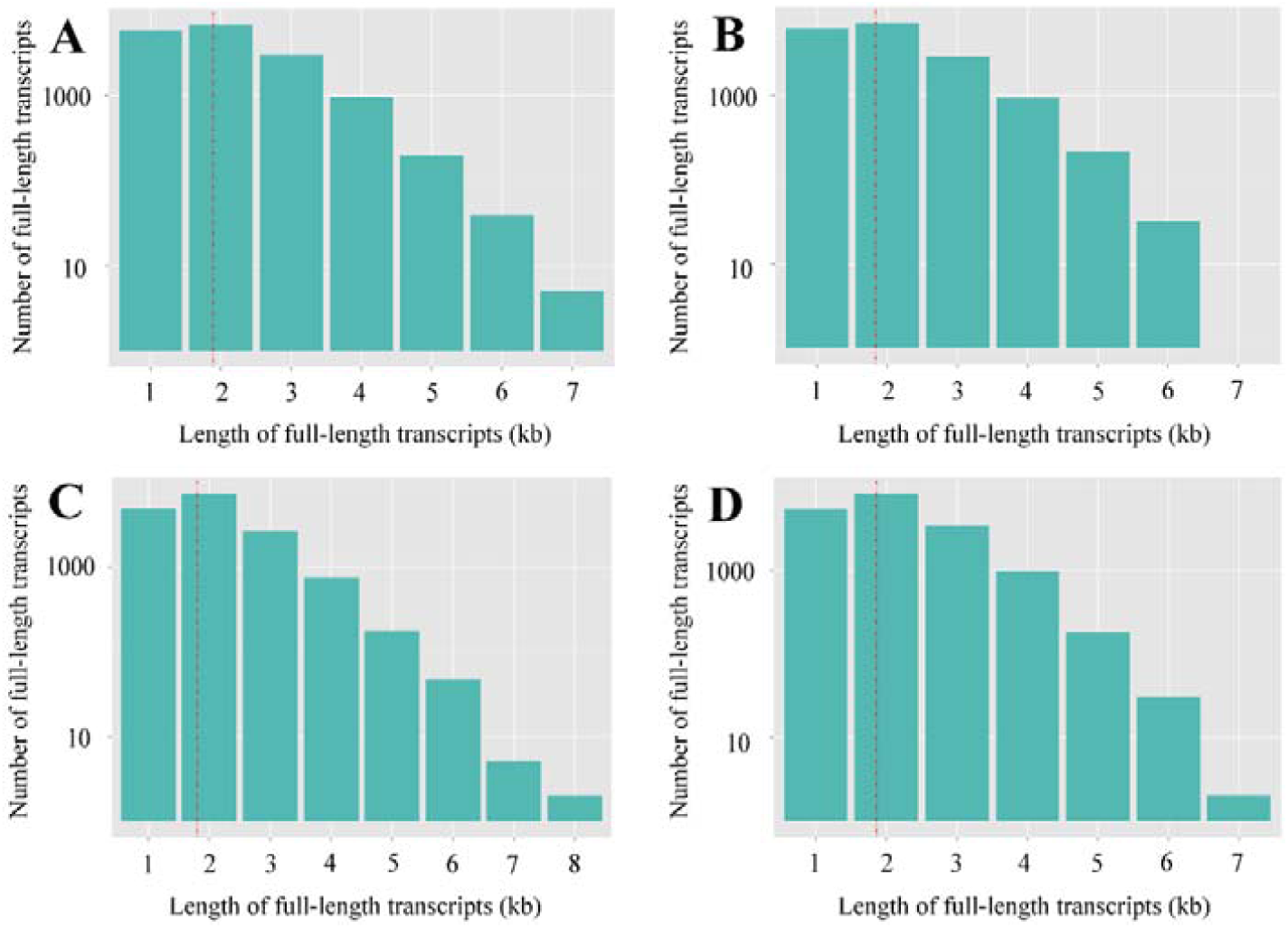
Length distribution of redundant reads-removed full-length transcripts from AmCK1 (A), AmCK2 (B), AmT1 (C) and AmT2 (D).

## Experimental Design, Materials, and Methods

### Honeybee midgut sample preparation

Spores of *N. ceranae* were previously prepared on basis of the method described by Cornman et al. [1] with some minor modifcations [2]. A bit of spores was detected via PCR detection and confirmed to be mono-specific [3] using previously developed specific primers [4]. The clean spores were immediately used for inoculation experiment.

Artificial inoculation of workers was conducted following our previous experimental protocol [2, 5]. The collected midgut samples were immediately frozen in liquid nitrogen and kept at −80 °C until sequencing.

### cDNA library construction and Nanopore sequencing

Firstly, total RNA of AmT1, AmT2, AmCK1 and AmCK2 were respectively extracted using TRizol reagent (Thermo Fisher, China) on dry ice, followed by reverse transcription with Maxima H Minus Reverse Transcriptase and concentration on a AMPure XP beads. Next, according to the manufacturer’s instructions (Oxford Nanopore Technologies Ltd, Oxford, UK), cDNA libraries were constructed from 50 ng RNA with a 14 cycles of PCR amplification using the cDNA-PCR Sequencing Kit (SQK-PCS109) and PCR Barcoding Kit (SQK-PBK004). Finally, the constructed cDNA libraries were sequenced using PromethION system (Oxford Nanopore Technologies Ltd, Oxford, UK) by Biomarker Technologies (China).

### Processing and quality control of Nanopore long reads

Raw reads were filtered with minimum average read quality score=7 and minimum read length=500 bp, and ribosomal RNA were removed after mapping to rRNA database. Next, full-length non-chemiric (FLNC) transcripts were determined by searching for primer at both ends of raw reads. Then, clusters of FLNC transcripts were obtained after mapping to the *Apis mellifera* genome (assembly Amel_4.5) with mimimap2 software [6]. Consensus isoforms were gained after polishing within each cluster with pinfish tools (https://github.com/nanoporetech/pinfish) and then mapped to the A. mellifera genome (assembly Amel_4.5) using minimap2. Ultimately, mapped reads were further collapsed using cDNA_Cupcake package (https://github.com/Magdoll/cDNA_Cupcake) with min-coverage=85% and min-identity=90%, and 5’ difference was not considered when collapsing redundant clean reads.

## Acknowledgments

This research was supported by the Earmarked Fund for China Agriculture Research System (No. CARS-44-KXJ7), the Science and Technology Planning Project of Fujian Province (No. 2018J05042), the Teaching and Scientific Research Fund of Education Department of Fujian Province (No. JAT170158), the Outstanding Scientific Research Manpower Fund of Fujian Agriculture and Forestry University (No. xjq201814), and the Scientific and Technical Innovation Fund of Fujian Agriculture and Forestry University (No. CXZX2017342, No. CXZX2017343).

